# Density-based optimization for unbiased, reproducible clustering applied to single molecule localization microscopy

**DOI:** 10.1101/2024.11.01.621498

**Authors:** Joseph L. Hammer, Alexander J. Devanny, Laura J. Kaufman

## Abstract

Single molecule localization microscopy (SMLM) has provided insight into the spatial organization of molecules at length scales below the diffraction limit of visible light. In SMLM data, density-based clustering approaches have proven to be valuable tools for probing the nanoscale structure of biological molecules, although little guidance is available for evaluating the accuracy of these results, which are often strongly dependent on user-input parameters. Here, we develop an efficient implementation of density-based cluster validation (DBCV) that can quantitatively evaluate clustering performance in SMLM-sized datasets without ground truth knowledge. We demonstrate that maximizing DBCV scores accurately identifies ground truth clustering in noisy, simulated datasets. By coupling DBCV score maximization with Bayesian optimization, we outline an optimization method, DBOpt, that selects unbiased input parameters for density-based clustering algorithms. We demonstrate that optimal input parameters can be selected for popular algorithms (DBSCAN, HDBSCAN, OPTICS) with minimal user input. Lastly, we show that DBOpt reports accurate feature sizes in 2D and 3D experimental datasets. Taken together, we propose an analysis pipeline that can be applied to a diverse array of experimental data that will improve the integrity and quality of cluster analyses in the broader scientific community.

## Introduction

Single molecule localization microscopy (SMLM) describes a class of super-resolution techniques commonly employed to overcome the diffraction limit associated with traditional microscopy.^1^ Many forms of SMLM exist, including stochastic optical reconstruction microscopy (STORM),^2^ photoactivatable localization microscopy (PALM),^3^ and point accumulation in nanoscale topography (PAINT).^4^ Regardless of chosen technique, SMLM data consists of point coordinates with associated uncertainties that correspond to fluorophore positions. While great advances have been made to develop new and refine established SMLM techniques, efficient and accurate analysis of the resulting data still presents major challenges.^5^ In coordinate-based SMLM data, clustering analyses are commonly performed to reveal insights into the complex underlying structure and spatial coordination of biological molecules.^5,6^ Among clustering methods, density-based clustering is a common choice for SMLM data, as it avoids biasing towards convex shapes like other clustering methods such as k-means, mean shift, and various hierarchical clustering algorithms.^7–10^ Instead, density-based methods allow arbitrary cluster shapes by identifying clusters as high density connected regions separated by regions of low density.^6,7,11–13^

A popular density-based clustering method is density-based spatial clustering of applications with noise (DBSCAN).^14^ DBSCAN connects points into clusters based on a reachability distance. DBSCAN is commonly employed for SMLM data and has been shown to perform well when ground truth information is available.^6,15^ A recent modification of DBSCAN, hierarchical DBSCAN (HDBSCAN), aims to improve DBSCAN by allowing clusters to vary in density through selection from a hierarchical tree constructed from mutual reachability distances.^16,17^ HDBSCAN is relatively understudied compared to DBSCAN, likely due to its more recent development. A third algorithm, ordering points to identify the clustering structure (OPTICS), allows variable density clusters to be identified through the ordered reachability of points.^18^ For each of these algorithms, at least two input parameters are required to identify clusters.

As with nearly all clustering algorithms, DBSCAN, HDBSCAN, and OPTICS present challenges in choosing input parameters.^12^ In each algorithm, parameter choice is not intuitive, even when domain knowledge is available. Thus, we suggest cluster validation should be implemented to guide parameter selection and increase reproducibility. Many choices for cluster validation exist, falling primarily into two classes, external and internal. External validation requires ground truth knowledge, allowing comparisons to be made between clustering algorithm outputs and the ground truth assignment of points into clusters.^19^ While effective, external validation is impractical in almost all real-world scenarios, where ground truth information is rarely available. Despite this, there are practical uses of external validation for experimental SMLM data. For example, Nieves et al. proposed a framework in which experimental data is compared to a set of simulated datasets to determine which simulation most closely matches the experimental data. A clustering algorithm and corresponding parameters are then chosen for experimental data that perform best on the nearest matching simulated data.^15^ To our knowledge, Nieves et al. provide the most comprehensive guidance for choosing clustering parameters for SMLM. However, critical limitations exist with this approach. The method is limited to the simulated datasets analyzed, restricting users to predefined structures and making the approach impractical for varied and/or complex clusters. Moreover, given that there can be no standard set of simulations for all possible clustering scenarios, outcomes could vary across research groups.

Alternatively, internal validation methods do not rely on ground truth information, but rather score the clustering based on intrinsic properties of the data.^19^ Many internal validation algorithms exist, with most unsuitable for validating non-globular clusters and therefore impractical for use with density-based clustering algorithms.^20,21^ Density based cluster validation (DBCV) is one of the few validation methods tailored to density-based algorithms,^21^ although it remains underutilized. In brief, DBCV evaluates clustered data by comparing the intra-cluster spread of points to the inter-cluster separation locally. Intuitively, clusters are scored higher as the spread of points within a cluster decreases and the separation between clusters increases. Unlike other validation methods, DBCV does not require input parameters, improving its strength as an unbiased evaluator.^21,22^

We propose a clustering optimization method that utilizes the internal validation metric DBCV to find optimal clustering input parameters. This method optimizes clustering by iteratively assessing the performance of different parameter combinations and selecting the combination with the highest validation scores. To our knowledge, the fastest publicly available implementations of DBCV remain far too slow for validating SMLM-sized datasets, especially when considering many distinct parameter combinations.^23–25^ Furthermore, naively sweeping parameters to find optimal clustering is not scalable to large parameter spaces. Therefore, this approach requires both a performance-efficient implementation of DBCV and a method to efficiently sweep large parameter spaces where there is little knowledge of appropriate parameter bounds.

Herein we (1) provide an efficient implementation of DBCV that is scalable to SMLM- sized datasets, (2) couple this improved DBCV implementation with Bayesian optimization to efficiently find DBCV maxima (DBOpt), and (3) demonstrate the efficacy of DBOpt by evaluating its performance on simulated and experimental datasets.

## Results

### DBCV Implementation

DBCV calculates individual cluster scores based on the intra-cluster sparseness and inter-cluster separation of each cluster (*Ci*) (**Eqn. 1**).^21^ An aggregate score (*DBCVscore*) summed over all clusters is computed from the individual cluster scores as a weighted average based on the number of points in each cluster (*NCi*) and the total number of points in the dataset, including noise (*Ntotal*) (see Methods, **Eqns. 2,3** for more details).^21^

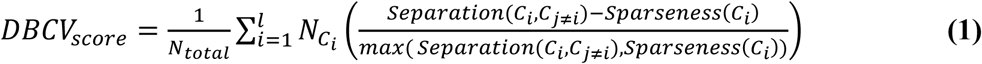

DBCV for SMLM must be scalable to large datasets. To our knowledge, the fastest implementations currently available are too slow for practical use on SMLM-sized data.^23–25^ To improve upon this, we quantified cluster separation leveraging a k-dimensional tree to find nearest neighbor distances of core points. A k-dimensional tree is constructed with an approximate time complexity *O*(*N*_*core*_(log(*N*_*core*_)) where *Ncore* are all clustered, core points as defined by DBCV.^26^ Once constructed, the separation value of each cluster can be calculated by querying the tree with approximate time complexity *O*(*N*_*C_i_ core*_(log(*N*_*C_i_ core*_ + 1)).^26^ With many other steps in the algorithm, the overall theoretical time complexity of the improved implementation (k-DBCV) is difficult to calculate. Thus, we benchmarked the performance of k-DBCV on simulated datasets, and show up to orders of magnitude increases in speed relative to previous implementations while maintaining efficient memory usage (**Fig. S1**).

### Density-Based Optimization

With an improved implementation of DBCV in hand, we now optimize clustering by selecting parameters that result in the highest DBCV score. DBOpt combines k-DBCV computation with Bayesian optimization to maximize the DBCV score of a clustering algorithm within a user-defined parameter space. Bayesian optimization provides an efficient method to maximize the output of a function without requiring exploration of every possible parameter combination, making it suitable for cases where there is little to no knowledge of optimal parameters and/or sensitivity to parameters.^27^ In brief, the optimization relies on a Gaussian prior function with an upper confidence bound acquisition function that balances maximization and exploration during optimization to iteratively select points for evaluation. Multiple iterations are performed to efficiently find the maxima^27,28^

The proposed cluster analysis pipeline is shown in **Scheme 1**. Before the optimization process, hyperparameters must be selected for DBOpt. This includes the lower and upper bounds of the parameters unique to the clustering algorithm that will be employed and the number of iterations to be performed. Recommended parameter bounds and number of Bayesian optimization iterations to perform are described in **Supporting Text 1**, **Fig. S2**.

**Scheme 1.**
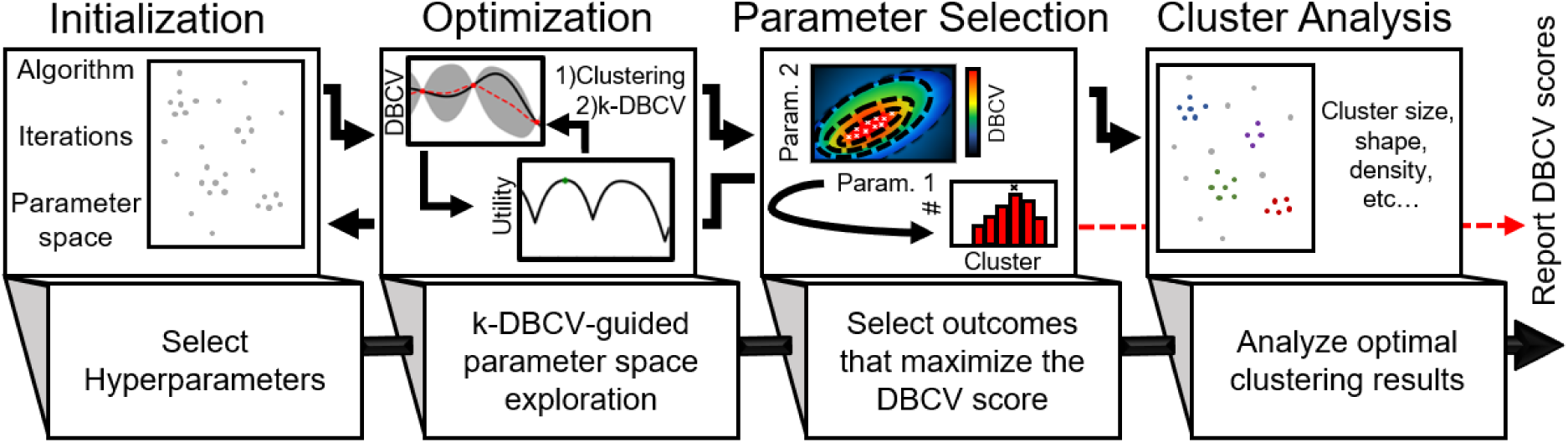
Schematic depiction of the cluster analysis pipeline. DBOpt is initialized with broad user hyperparameter selections, the parameter space is explored, and the parameters identifying clusters with maximum DBCV score are selected for clustering. Following DBOpt, cluster analysis is performed and DBCV scores are reported.

The optimization step produces a series of parameter combinations with corresponding DBCV scores between -1 and 1, with -1 automatically assigned when there are fewer than two clusters identified. The output commonly contains several parameter combinations that produce equal maximum aggregate DBCV scores (to within two significant figures). DBOpt selects between these by choosing parameters for which median individual cluster scores are highest.

Additional DBOpt runs should be performed in cases where the initial hyperparameters for parameter bounds may be too small or too large to effectively optimize the parameters. Evidence of insufficient size would be maximum DBCV scores at the edge of the parameter space. Evidence of an optimization space that is too large would be a small number of parameters evaluated near the DBOpt selected parameters or a small number of scores near the maximum DBCV score. Confirmation of proper optimization can be achieved by performing DBOpt multiple times on the same dataset from a random set of initial parameters and ensuring the maximum score converges to approximately the same value. The resulting clusters from DBOpt will have both a global validation score and individual cluster scores, allowing for comparisons across datasets as well as outlier detection within datasets, respectively. After DBOpt, cluster analysis can be performed and the selected clustering parameters along with the cluster validation scores should be reported.

### Validation of DBOpt with Simulated Data

To assess DBCV as a validation metric and test the ability of DBOpt to identify optimal parameters, we evaluated simulated datasets. We simulated SMLM data across five categories of clusters, with representative datasets shown in **Fig. 1**. For each category of clusters, twenty-five unique datasets on an ≈ 3 x 3 μm 2D plane were simulated as described in Methods. The simulations varied cluster size, shape, density, number of clusters, and noise density (**Supporting Text 2**, **Table S1**).

**Fig 1.**
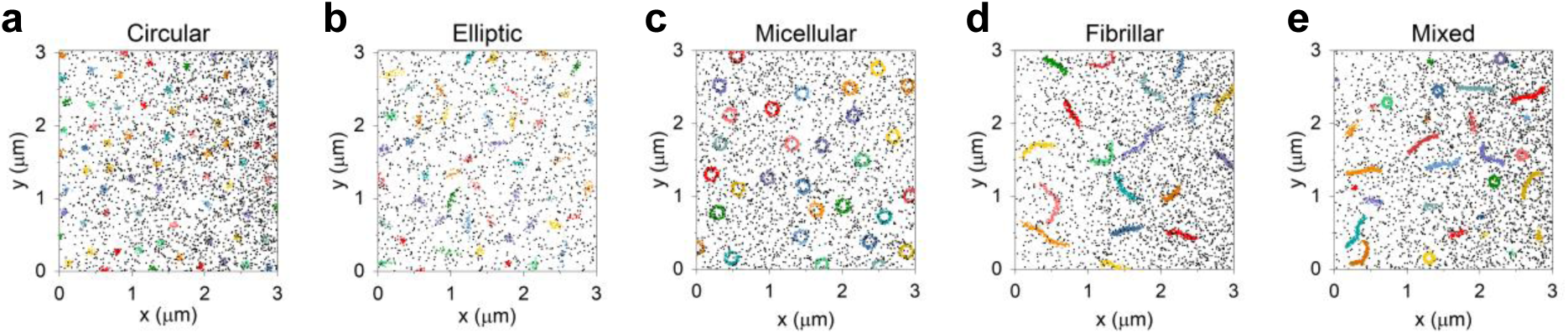
Representative plots of simulated **a)** circular (C02), **b)** elliptic (E14), **c)** micellular (M22), **d)** fibrillar (F18), and **e)** mixed (V12) cluster scenarios. Alphanumeric code in parentheses corresponds to simulations described in Table S1. Simulated clusters are shown in color, noise is shown in grey. **a, d,** and **e** have gradient noise. **b** and **c** have homogeneous noise.

To first evaluate the performance of each clustering algorithm, we systematically explored parameter combinations and evaluated their performance against ground truth information with the external validation V-measure (**Eqn. 4**).^29^ We refer to this method as a naive external validation sweep (EVS). For all simulations, naive EVS was performed with DBSCAN, HDBSCAN, and OPTICS as described in Methods. Separately, DBOpt was performed to assess its ability to match EVS performance in the absence of ground truth information. Hyperparameter selection for DBOpt is described in Methods. Representative results from EVS and DBOpt on challenging simulations (> 50% noise) are shown in **Fig. 2**. The contour plots associated with EVS and DBOpt show similar character. We plotted the resulting clusters from the highest scoring parameters for each clustering algorithm based on naive EVS V-measure scores and, separately, DBOpt DBCV scores (**Fig. S3**). Between all algorithms, we show the highest scoring cluster outcome in **Fig. 2**. Naive EVS identifies clusters that qualitatively match ground truth information for DBSCAN while HDBSCAN and OPTICS deviate from ground truth for more complex clustering scenarios, indicating a weaker performance by these algorithms (**Fig. S3**).

**Fig 2.**
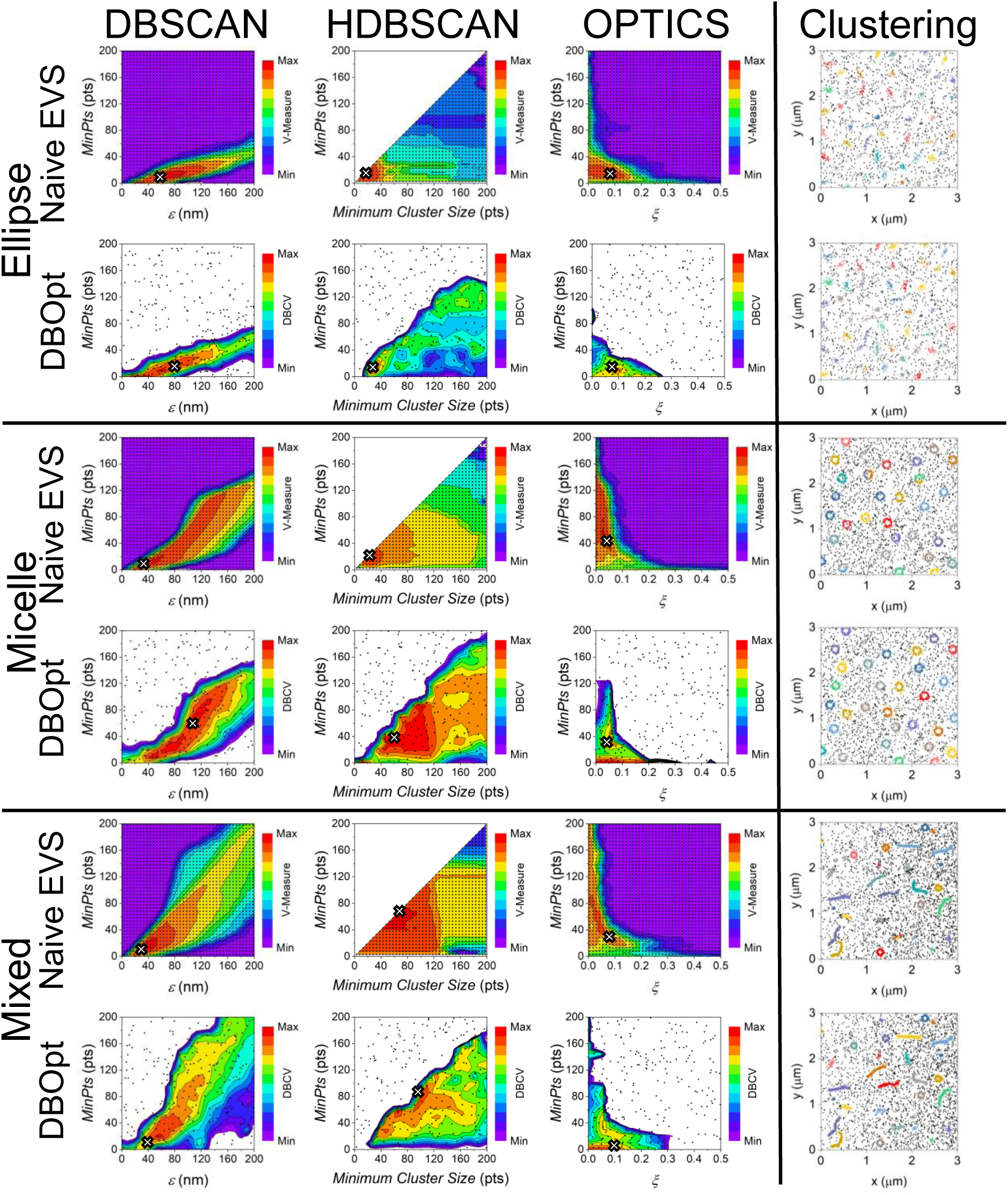
Naive EVS and DBOpt parameter sweeps of elliptic (E14), micellular (M22), and mixed (V12) cluster scenarios for DBSCAN, HDBSCAN, and OPTICS, with optimal EVS and DBOpt cluster outcomes plotted in the right panel. Parenthetical notations indicate the simulated data as described in Table S1. Small points represent each parameter combination that is scored from which the contour plot is prepared; white “X” represents the optimal chosen parameters. Right panel plots correspond to those marked with * in Fig. S3.

When EVS-identified clusters agreed with ground truth, DBOpt qualitatively matched naive EVS performance (**Fig. S3**). To compare the quantitative performance of DBOpt on every simulated dataset, we calculated the V-measure scores of the parameters chosen by DBOpt and plotted them vs. the maximum V-measure scores achieved by naive EVS for each algorithm (**Fig. 3a-c**). Here, DBOpt paired with DBSCAN most closely matches scores achieved with naive EVS, with weaker performance for some fibrillar and mixed clustering scenarios. Also shown is the combined performance, where the selected parameters correspond to the maximum DBOpt DBCV score between all clustering algorithms (**Fig. 3d**).

**Fig. 3.**
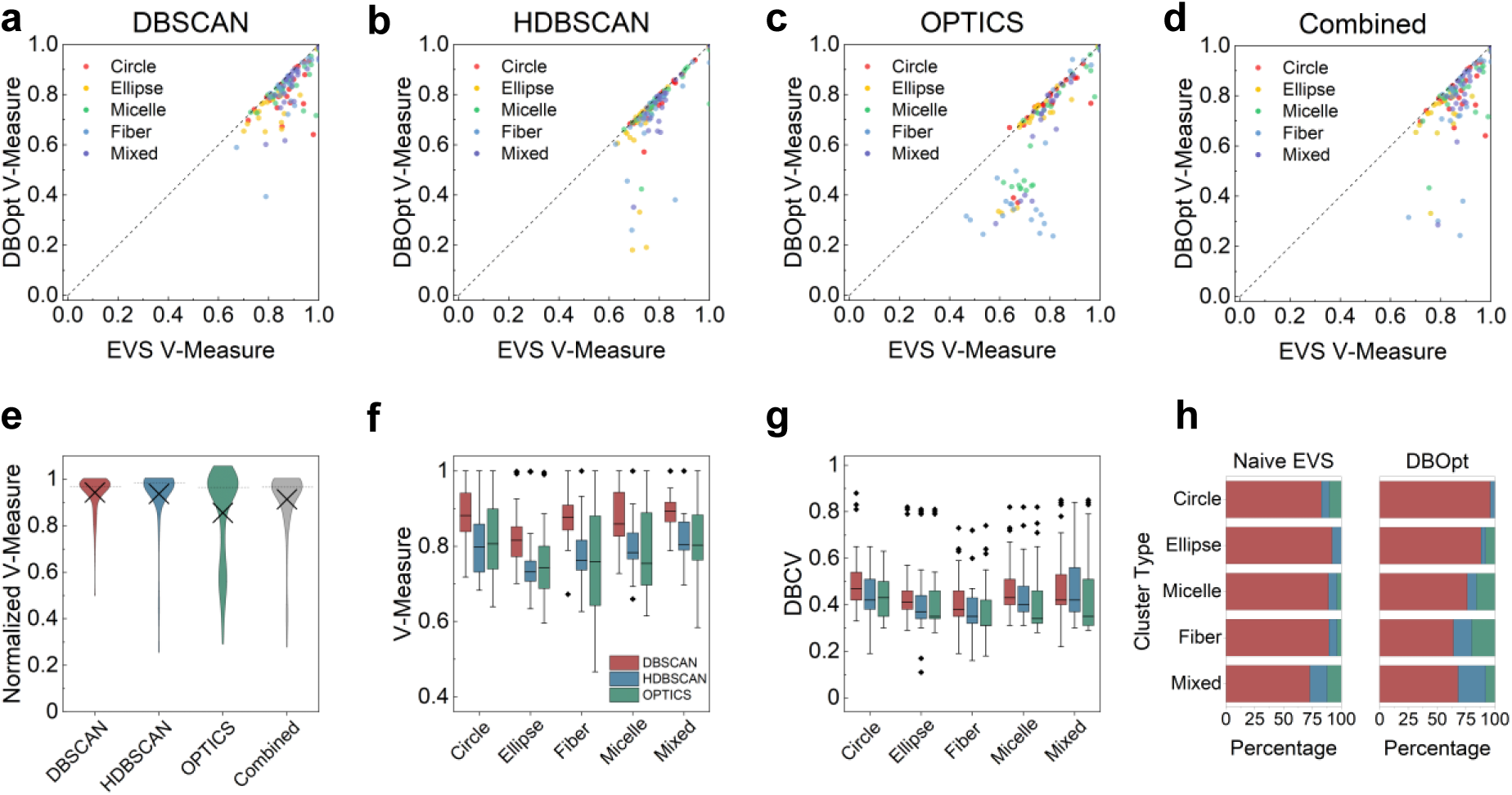
Comparisons between maximum V-measure scores from naive EVS and V-measure scores from DBOpt- selected parameters for **a)** DBSCAN, **b)** HDBSCAN, **c)** OPTICS, and **d)** all algorithms combined, where the DBOpt parameters are selected from the maximum DBCV score across all algorithms. **e)** Distributions of DBOpt V-measure scores to maximum naive EVS V-measure scores (dotted line represents median and “X” represents mean). **f)** Maximum V-measure and **g)** maximum DBCV score comparisons between algorithms for each cluster simulation type. Box indicates 25^th^ and 75^th^ percentile, whiskers indicate 5^th^ and 95^th^ percentile, the interior line indicates the median score, and outliers are shown as points. **h)** Percentage of the simulations for which each algorithm had the highest V-measure and DBCV scores, respectively. The legend in **f** applies to **f-h**.

The overall performance of DBOpt relative to naive EVS is shown by taking the ratio of the V-measure scores, with distributions shown in **Fig. 3e**. The median normalized V-measure was 0.97, 0.98, 0.97, and 0.97 for DBSCAN, HDBSCAN, OPTICS, and all algorithms combined, respectively, indicating that in most cases DBOpt effectively identified clusters as if ground truth assignments were known. To broaden the simulations to capture more experimental scenarios, we also evaluated DBOpt performance on multi-emitter data. Here, each originally simulated point may have multiple corresponding points drawn from the same uncertainty distribution, resulting in more localizations for both clustered and noise points (**Supporting Text 3, Fig. S4**). DBOpt remains robust in most multi-emitter scenarios for DBSCAN and HDBSCAN. Furthermore, when accounting for fluorophore to antibody ratios when setting the *MinPts* hyperparameter, DBOpt fully recovers and in some cases improves its performance in the presence of multi-emitters (**Fig. S4c**). Regardless of parameter bounds, OPTICS performance is diminished by the presence of multi-emitters, limiting its practicality for evaluating SMLM data.

We further compared V-measure and, separately, DBCV scores of optimal parameters between algorithms for every simulated SMLM dataset (**Fig. 3f-h**). These comparisons indicate that DBSCAN generally scores higher than HDBSCAN and OPTICS for each cluster type. In cases where noise was low, HDBSCAN occasionally returned a slightly higher V-measure score and in rare circumstances, OPTICS returned the highest DBCV score. Finally, we compared the runtime of each algorithm with DBOpt. Here, DBOpt paired with DBSCAN and HDBSCAN had similar runtimes, while OPTICS was slower for every simulation (**Fig. S5**).

We note that, as can be appreciated from **Fig. 3g**, DBCV scores are relatively low compared to the theoretical maximum of 1 as well as to the maximum V-measure scores. This is primarily because DBCV values are reduced by the presence of noise (**Eqn. 1**). With simulated datasets, we can ensure that clusters are properly identified even when DBCV scores are low because ground truth information is available. However, for experimental data, a threshold may exist where scores are too low to distinguish accurate clustering from the clustering of noise. To determine this threshold, we evaluated DBOpt on simulated 2D noise and show the threshold decreases as the lower bound of the *MinPts* parameter increases (**Supporting Text 4, Fig. S6a**).

Extension of traditional SMLM to 3D imaging has revealed complex cellular structures in unprecedented detail.^30,31^ To test whether DBOpt performs well on 3D data, ten unique datasets containing 3D ellipsoidal clusters were simulated (**Table S2**) as described in Methods. Naive EVS and DBOpt were performed separately for each algorithm. A representative simulation is shown in **Fig. 4a**, with the clustered data resulting from DBOpt shown in **Fig. 4b**. Comparisons between DBOpt and external validation of each algorithm is shown in **Fig 4c-f**. The normalized V-measure scores for DBSCAN, HDBSCAN, and OPTICS were 0.98, 0.99, and 0.98, respectively. Comparing across algorithms revealed that DBSCAN scores higher than HDBSCAN and OPTICS for every simulation with both naive EVS and DBOpt. Simulated noise was evaluated to assess the presence of a threshold in 3D under which cluster identification cannot be distinguished from clustering noise (**Supporting Text 4, Fig. S6b**). Overall, the noise threshold was reduced by the increase in *MinPts* and dimensionality.

**Fig. 4.**
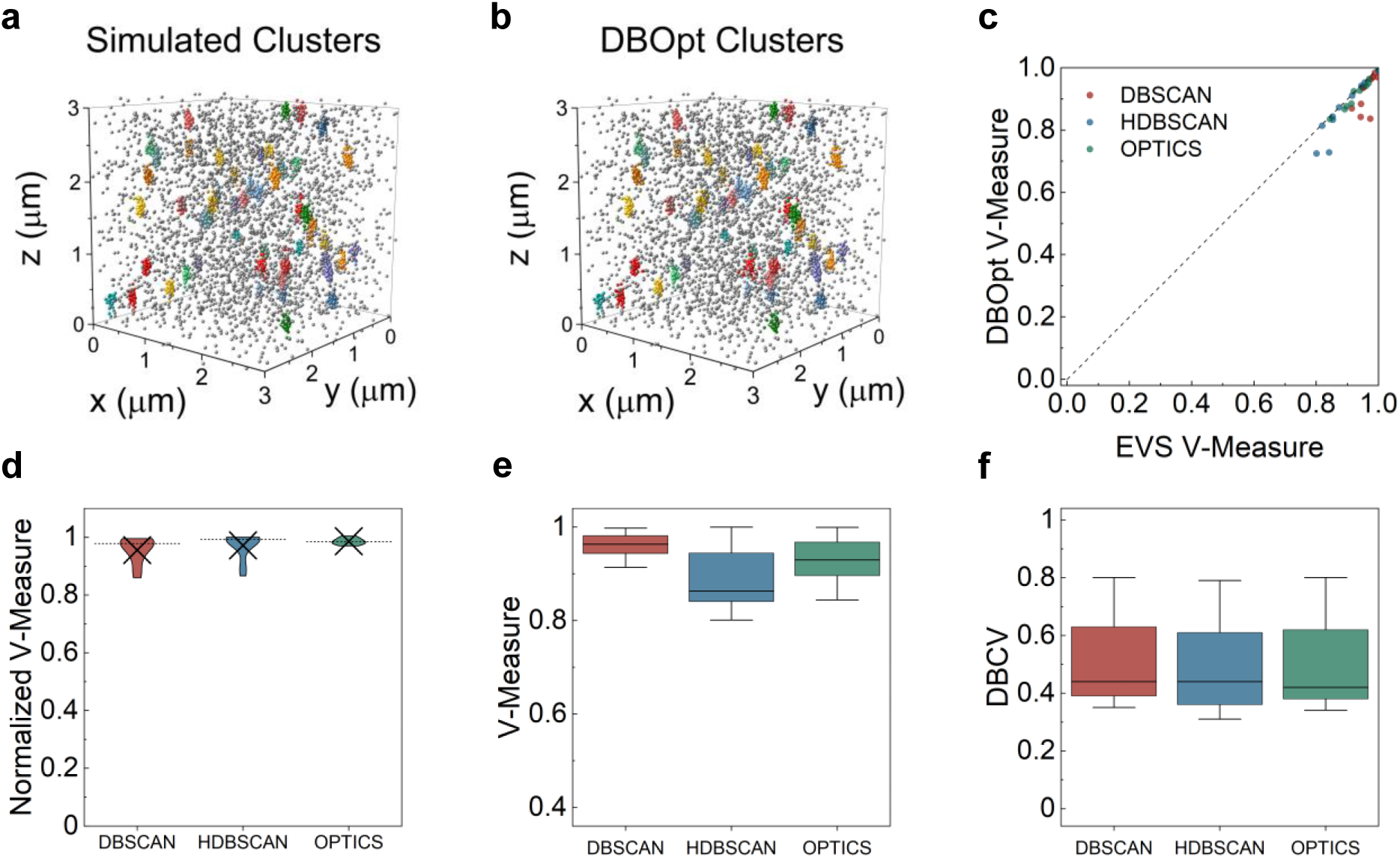
a) Representative simulation (S01, Table S2) of 3D clusters, **b)** optimal clustering of the representative simulated data via DBOpt, **c)** comparisons of V-measure scores between naive EVS and V-measure scores for parameters chosen with DBOpt, **d)** distributions of the ratio of DBOpt V-measure scores to maximum naïve EVS V-measure scores (dotted line represents median and “X” represents mean), and **e)** optimal V-measure and **f)** DBCV scores for each algorithm found via naive EVS and DBOpt, respectively. Box indicates 25^th^ and 75^th^ percentile, whiskers indicate 5^th^ and 95^th^ percentile, and the interior line indicates median.

### Experimental SMLM Analysis

To demonstrate a practical use case for DBOpt, we quantified the size and shape of β1 integrin nanoclusters in the ligand-bound conformation within focal adhesions. MBA-MB-231 cells were fixed and regions of interest (ROI) were identified by the presence of vinculin, an adapter protein localized to focal adhesions.^32^ β1 integrin was labeled with an antibody specific to the ligand bound conformation.^33^ A widefield image of cells labeled for vinculin and β1 integrin was acquired, followed by 2D STORM imaging of β1 integrin. A representative reconstructed image is shown in **Fig. 5a-b**. To identify clusters, DBOpt coupled with DBSCAN was performed on each acquired dataset (**Fig. 5c**). The optimal parameters identified with corresponding DBCV scores are shown in **Table S3**. A representation of resulting clusters, colored to depict individual cluster scores, is shown in **Fig. 5d**. For analysis, clustered integrin localizations within ROI co-localized with vinculin were selected. Identified clusters with positive individual cluster scores were analyzed (**Fig. 5e**). Cluster shape (aspect ratio) and size were determined (**Fig. 5f**) as described in Methods. Our results indicate a short-axis median FWHM of 53 nm and a median aspect ratio of 1.51. These results are in close agreement with previously reported integrin cluster sizes.^34,35^

**Fig 5.**
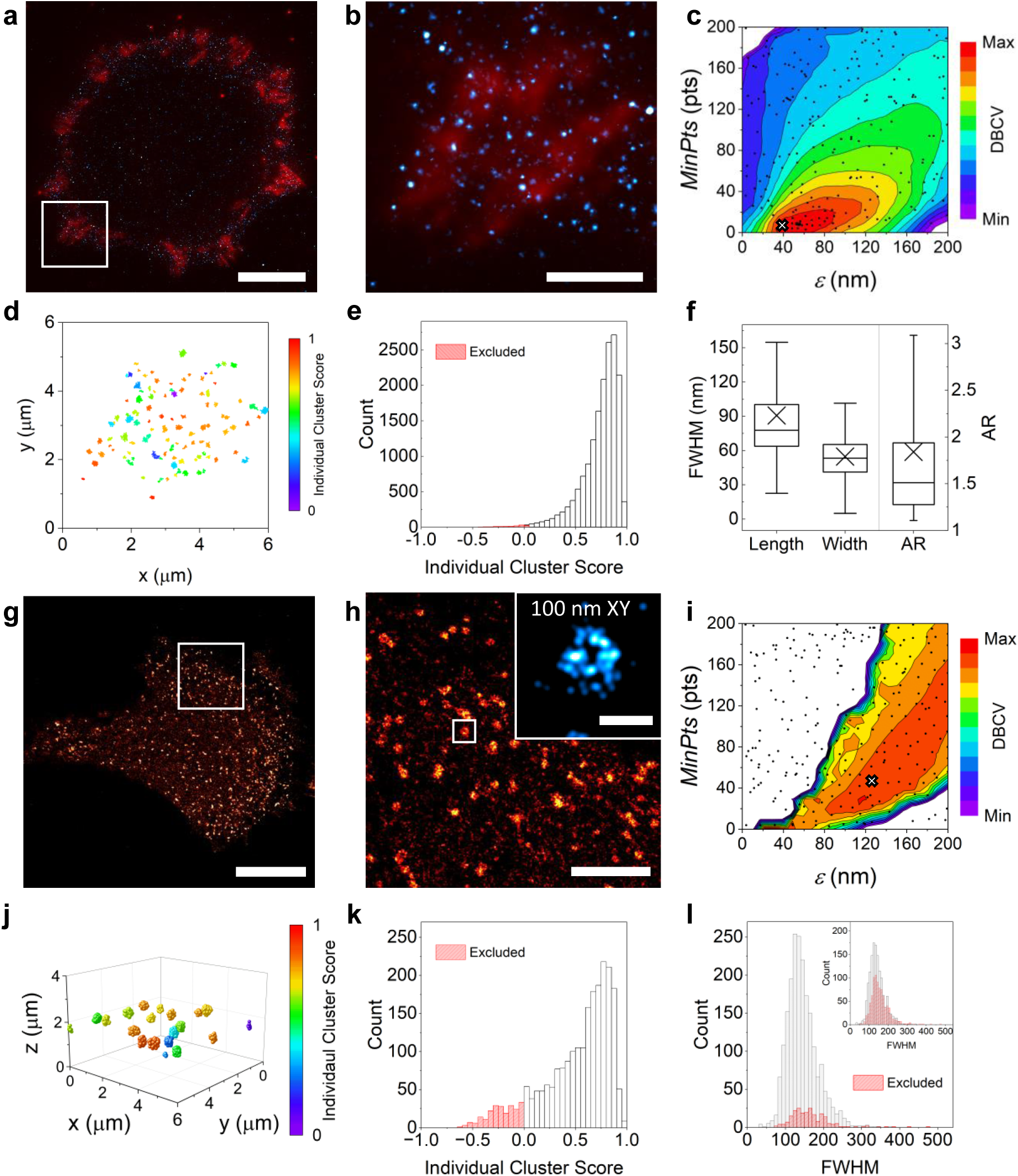
a) STORM image (Integrin 02, Table S3) of a cell co-labeled for vinculin (red) and active β1 integrin (cyan). (scale bar: 10 μm) **b)** Selected region of interest (white box in (**a**); scale bar: 2 μm). **c)** DBOpt parameter sweep of integrin (Integrin 02) localizations with DBSCAN. **d)** Selected region (corresponding to (**b**)) of identified integrin clusters from DBOpt; color depicts individual cluster score. **e)** Distribution of individual cluster scores. All clusters with an individual cluster score ≥ 0 were analyzed. **f)** Cluster width, length, and aspect ratio (AR). **g)** STORM image (Clathrin 01, Table S3) of a cell labeled for clathrin (red) (scale bar: 10 μm). **h)** Selected region (white box in (**g**); scale bar: 2 μm)) with inset showing a 100 nm axial region of a selected clathrin cluster projected in 2D (scale bar: 0.2 μm). **i)** DBOpt parameter sweep of clathrin (Clathrin 01) localization with DBSCAN. **j)** Selected region of identified clathrin clusters from DBOpt; color depicts individual cluster score. **k)** Distribution of individual cluster scores; clusters scoring < 0 were excluded. **l)** Distribution of FWHMs of identified clathrin clusters, inset depicts the distribution with individual cluster scores < 0.5 excluded.

To analyze a more challenging dataset, we sought to quantify clathrin-coated pits in MDA-MB-231 cells. To do this, we performed 3D STORM on fixed cells labeled for clathrin. A representative reconstruction of all molecules projected in 2D is shown in **Fig. 5g**, with a select region shown in **Fig. 5h**, illustrating the pit structure. DBOpt was performed on each 3D dataset with DBSCAN (**Fig. 5i**). Maximum DBCV scores were between 0.09 and 0.15, with the *MinPts* parameter ranging from 34 to 112, above the corresponding noise threshold (**Supporting Text 5**, **Table S3**). A selection of identified clusters is shown in **Fig. 5j**. When evaluating clusters with a positive individual cluster score, the mean FWHM of clathrin clusters was found to be 145 ± 40 nm, in agreement with previously reported results (**Fig. 5l**).^36^ Excluding clusters with low individual cluster scores may be useful in particular cases. Here, evaluating clusters with a stricter individual cluster score requirement (individual cluster score ≥ 0.5; inset **Fig. 5l**) did not affect results (FWHM 141 ± 39 nm). DBCV scores and chosen parameters for all SMLM datasets are provided in **Table S3**.

## Discussion

Cluster analysis is a common and important step in interpreting SMLM data. However, the importance of proper parameter selection is often overlooked, as evidenced by the relative lack of guidance for parameter selection in the literature. For these reasons, we (1) developed an efficient implementation of the internal validation metric DBCV, termed k-DBCV for its use of a k-dimensional tree, and (2) designed a procedure for leveraging k-DBCV to choose optimal clustering parameters, incorporating Bayesian optimization to efficiently optimize large parameter spaces. Taken together, the DBOpt method provides a valuable tool for selecting robust and reproducible clustering parameters without the need for domain knowledge.

The results from simulated datasets suggest that in most scenarios DBOpt, without ground truth information, is nearly equal in performance to naive EVS as evaluated via V-measure (**Fig. 3**). Furthermore, we demonstrate DBOpt on experimental data and show sensible results from cluster evaluation. Among the clustering algorithms employed, DBSCAN was the best performing algorithm for most simulated datasets. Paired with its relative simplicity and speed, we recommend DBSCAN when clustering SMLM data. While we evaluated DBSCAN, HDBSCAN, and OPTICS, DBOpt can be readily adapted to evaluate performance against any density-based algorithm.

Through the approach outlined herein, we expect that DBOpt will improve both the integrity and reproducibility of SMLM clustering. While this work highlights the utility of DBOpt for SMLM data, the importance of accurate clustering extends across biology and into many other fields of study.^37–40^ Thus, we expect DBOpt to have many practical use cases outside of SMLM.

## Methods

### Simulations

Simulated data was generated using our custom-built Python library (ClustSim: https://github.com/Kaufman-Lab-Columbia/ClustSim). Generally, for each cluster a centroid was randomly chosen on the ≈ 3 x 3 μm simulation plane and points were placed around this centroid. When simulating in 3D, an additional axial dimension of 3 μm was added. Circular, spherical, and elliptic clusters were built by making random selections from a normal distribution centered around the centroid in each dimension. The cluster width was defined as 4 standard deviations of the underlying normal distribution. For elliptic clusters, the distribution was stretched randomly along one direction, defined by the desired aspect ratio. For micellular clusters, points were randomly and uniformly distributed between an inner and outer diameter with the inner diameter defined as two-thirds of the outer diameter. Fibrillar clusters were generated via a three-step process. First, the fiber backbone was grown from a random starting point along a simulated trajectory, defined by an angular path dictating the direction of longitudinal growth. The angular path was generated using the method described by Bi et al.^41^ Subsequently, lateral point deposition to a specified density was conducted around each backbone point using a normal distribution, with 4 standard deviations of the distribution equal to the reported widths. Finally, for clusters containing a variety of shapes, clusters were simulated separately and merged, ensuring clusters were adequately separated by eye.

Each simulation was generated on a simulation plane containing either randomly distributed or gradient noise to mimic inhomogeneous illumination. Gradient noise was generated by increasing the percentage of points distributed every 300 nm in the x direction such that the right-most side of the simulation had approximately four times more noise points than the left-most side. For multi-emitter simulations, the number of localizations at each target point was drawn from a Poisson distribution with a mean of 3 positions per molecule.

In all cases, after placement, point positions were relocated in the x, y, and (where relevant) z directions to mimic uncertainties associated with SMLM imaging. Each molecule was moved within a FWHM defined from a log-normal distribution.^42^ This log-normal distribution was set with a mean uncertainty of 20 nm for the lateral directions and 50 nm axially for 3D simulations with a standard deviation of 5.7.^30,42,43^ Parameter information for each simulation can be found in **Supporting Text 2** and **Tables S1-2**.

### DBOpt

DBOpt (DBOpt: https://github.com/Kaufman-Lab-Columbia/DBOpt) was performed by combining the improved implementation of DBCV (k-DBCV: https://github.com/Kaufman-Lab-Columbia/k-DBCV) with Bayesian optimization. Bayesian optimization was performed with a pre-built library using a Gaussian prior with an upper confidence bound acquisition function (BayesianOptimization: https://github.com/bayesian-optimization/BayesianOptimization).^27^ For all simulations, hyperparameters were chosen to attempt to cover the relevant parameter space for all clustering scenarios and simulations while also testing the data against the minimum possible *MinPts* parameters for k-DBCV. For all simulations, 40 random sets of parameters were probed initially followed by 200 optimization iterations (**Supporting Text 1**). The parameter space explored was 3 to 200 for all parameters of DBSCAN and HDBSCAN and the *MinPts* parameter of OPTICS. For OPTICS, the *ξ* parameter space was optimized between 0.005 and 0.5.

At each optimization iteration, the DBCV score was calculated as described in **Eqn. 1**.^21^ Here, sparseness and separation are defined as the largest intra-cluster and smallest inter-cluster mutual reachability distances (*MRD*) between nearest neighboring core points, respectively. The mutual reachability distance between points is calculated as:

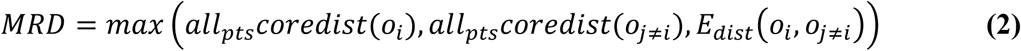

Here, *oi* is a point within the dataset, *oj* is any other point, *Edist* is the Euclidean distance between points, and *allptscoredist* is defined as:

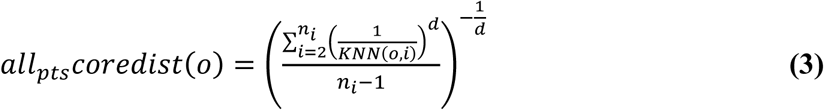

where *d* is the number of dimensions, *ni* is the points in the cluster *i*, and *KNN* is the Kth nearest neighbor from point *o* in cluster *i*. We note that a minimum of three points is required for a cluster to have at least one core point. Here, we require core points to compute individual cluster scores; therefore k-DBCV prohibits clusters with fewer than three points and automatically reclassifies points belonging to these clusters as noise After completing the optimization iterations, the parameter combinations with the highest DBCV scores to two significant figures were selected. Subsequently, from this set, the single parameter combination with the highest median individual cluster scores was chosen as the optimal clustering assignment. The data was then clustered with those parameters, and in the case of simulated data, external validation was performed for comparison to ground truth information and naive EVS.

### External Validation

Naive EVS was performed by analyzing every fifth value of each parameter between 3 and 200 for DBSCAN, HDBSCAN, and the *MinPts* OPTICS parameter. The *ξ* parameter of OPTICS was analyzed at every 0.0125 between 0.005 and 0.5. V-measure was employed for external validation. V-measure is given by:

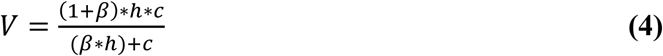

Here, homogeneity (h) and completeness (c) were calculated for comparison to ground truth information as described by Rosenberg et al.^29^ and β was set to 1 to weight contributions equally.

### Cell Preparation

MDA-MB-231 cells obtained from American Type Culture Collection were used for both integrin and clathrin experiments. Cells were cultured at 37 °C and 5% carbon dioxide in high glucose DMEM (Fisher Scientific) with 10% (v/v) fetal bovine serum (Gibco), 1% (v/v) 100x penicillin-streptomycin-amphotericin B (MP Biomedicals), and 1% (v/v) 100x nonessential amino acid solution (Gibco). Prior to cell experiments, 35 mm high tolerance dishes (P35G-0.170-14-C: MatTek Corporation) were coated with 1 mL of 50 μg/mL acid solubilized rat tail collagen type I (Advanced Biomatrix) in sterile filtered 20 mM acetic acid (97+%, Sigma Aldrich) for one hour. The plates were washed three times with 1X phosphate buffered saline (PBS) (Cytiva). Tetraspeck microspheres (Invitrogen) of diameter 0.1 μm were added at a concentration of approximately 4*10^15^ microspheres/mL in 1X PBS for 10 minutes at room temperature to deposit fiducials for later drift correction in post-processing. The plates were washed again three times with 1X PBS.

Cells were detached with Accutase (MP Biomedicals) and then seeded onto the coated, high tolerance 35 mm dishes at a density of 100,000 cells/dish in 2 mL high glucose DMEM and incubated for 1.5 hrs before fixing. Prior to fixing, cells were washed twice with 37°C 1X PBS. For β1 integrin labeling, cells were fixed by initially adding 0.3% glutaraldehyde (8% EM grade solution, Electron Microscopy Sciences) and 0.25% Triton-X 100 (10% solution, EMD Millipore Chemicals) in 1X PBS at 37 °C for 1 min followed by 4% methanol free paraformaldehyde (16% solution, Thermo Scientific) in 1X PBS at 37°C for 10 min. For clathrin experiments, 4% methanol free paraformaldehyde in 1X PBS was added for 10 min at 37 °C. After fixing, quenching with 50 mM ammonium chloride (Sigma Aldrich) for 15 min was performed. Cells were washed for 10 minutes three times with 1X PBS. Triton X-100 (0.2%) was applied for 10 min to permeabilize the cell membrane, followed by three 10 min washes with 1X PBS.

Active β1 integrin was labeled with 9EG7 monoclonal antibody (BD Biosciences). The antibody was first conjugated to Alexa Fluor 647 NHS ester (Invitrogen). Alexa Fluor 647 NHS ester was dissolved in anhydrous dimethyl sulfoxide (DMSO) (Sigma Aldrich) and dried for storage. The aliquots were desiccated and stored at -20°C. Before conjugation, bovine serum albumin (BSA) was removed (when applicable) from the stock antibody solution with an antibody conjugation kit according to manufacturer instructions (Abcam). After BSA removal, the antibody concentration was measured via UV-vis with a Nanodrop Spectrophotometer (Thermo Scientific). The antibody was then conjugated with Alexa Fluor 647 NHS ester at a 4:1 fluorophore to antibody molar ratio for 30 minutes by adding 2 μL of reconstituted Alexa Fluor 647 in sterile DMSO (Sigma Aldrich) to 10 μL of 0.5 M sodium bicarbonate (7.5% stock, Gibco) and 40 μL of antibody solution in 1X PBS. Following conjugation, the antibody was purified with an Antibody Conjugate Purification Kit (Invitrogen). Briefly, the column was rinsed three times with 1X PBS and centrifuged at 1100 x g. The antibody was then added to the column and incubated for 5 minutes at room temperature. The final solution was collected via centrifugation at 1100 x g. The resulting fluorophore to antibody ratio was measured via UV-vis and calculated to be 1.6:1.

The conjugated 9EG7-Alexa Fluor 647 was diluted in 1% BSA (w/w) (Fisher Bioreagents) in 1X PBS for a final concentration of 10 μg/mL along with 10 μg/mL EPR8185 anti-vinculin Alexa Fluor 488 antibody (Abcam). To label cells, 100 μL of antibody solution was added to the plate to fully cover the inner surface of the dish. The solution was incubated for 18 hrs at ∼ 4 °C. After labeling, the cells were washed three times with 1% BSA in 1X PBS.

To label clathrin, a polyclonal anti-clathrin heavy chain antibody (Abcam) was diluted to a final concentration of 3.3 μg/mL with 1% BSA (w/w) in 1X PBS. 100 μL of antibody solution was added to the plate to fully cover the surface and the solution was incubated for 18 hrs at ∼4 °C. The plate was washed three times with 1% BSA (w/w) in 1X PBS for 10 min. 100 μL of 4 μg/mL secondary, goat anti-rabbit Alexa Fluor 647 antibody (highly cross-adsorbed, Invitrogen) in 1% BSA (w/w) in 1X PBS was then added and incubated at room temperature for one hour. The dish was washed again three times with 1% BSA (w/w) in 1X PBS following secondary antibody labeling.

### Imaging

Prior to imaging, 1 mL of freshly prepared OxEA imaging buffer was added to the samples.^44^ The buffer was composed of 3% (v/v) Oxyfluor (Oxyrase), 20% (v/v) sodium DL lactate (60% stock, Sigma Aldrich), and 50 mM cysteamine hydrochloride (Sigma Aldrich), all in 1X PBS with pH adjusted to 8-8.5 with 1 N NaOH (Sigma Aldrich). Images were acquired on a Zeiss Elyra 7 microscope. For 2D experiments, an initial image of vinculin was acquired using a 488 nm laser via total internal reflection fluorescence (TIRF). For 2D SMLM, localizations were acquired over 30,000 frames with an exposure time of 30 ms in a TIRF configuration. For 3D acquisition, the microscope relies on a spatial light modulator to split the vertically polarized light into two lobes, forming a double helix point spread function (DH-PSF).^45^ Prior to imaging, a 0.1 μm Tetraspeck microsphere was imaged for calibration of the DH-PSF according to manufacturer instructions such that z-position of the PSFs could be extracted after acquisition. The localizations were acquired over 50,000 frames with an exposure time of 30 ms via highly inclined and laminated optical sheet (HILO) microscopy.

To process localizations found in 2D, the ThunderSTORM plugin was used within ImageJ following the recommended protocol in the ThunderSTORM user guide.^46^ Images were first filtered using a wavelet filter (B-spline order of 3 and B-spline scale of 2), after which molecules were identified using the local maximum approach (8 connected neighbors) with a peak intensity threshold of 1.5 times the standard deviation of the first wavelet level. Identified molecules were fit to an integrated Gaussian using the maximum likelihood method, with a fitting radius of 5 pixels and an initial standard deviation of 1.6 pixels. Spurious localizations were removed from reconstructed images by applying a minimum intensity cutoff of 50 photons, restricting the standard deviation of the Gaussian fit over the emission peak to 50 – 250 nm, and removing molecules with a lateral localization uncertainty greater than 35 nm. Lateral stage drift was corrected by tracking positions of approximately 3-6 fiducial markers (Tetraspeck microspheres, see above plating procedure) during the course of image acquisition. Axial drift was limited by using the microscope autofocus (Zeiss, Definite Focus system) in combination with a piezo stage to continuously maintain axial position for the duration of the imaging experiment. After drift correction using ThunderSTORM, molecules within 20 nm of each other were merged when in the on-state consecutively between frames with a maximum off time tolerance of three frames. Processing of 3D STORM data was performed within the Zen Black software (Zeiss). Axial position was first determined from the DH-PSF calibration. To remove outliers, the lateral localization uncertainty was filtered to be between 5 and 35 nm and the axial uncertainty was filtered to be between 5 and 60 nm. The background variance of the number of photons was set to less than 80. The images were drift corrected both laterally and axially with the 0.1 μm Tetraspeck microsphere fiducials.

For 2D integrin images, DBOpt was performed on the full cell. After clustering with the optimal parameters found for DBSCAN, clusters that fell within ROIs defined by the presence of vinculin were analyzed. For clathrin localization, DBOpt was first run on the full cell in 3D and identified clusters were projected onto an x-y plane. The covariance matrix of each cluster was used to find the long-axis (length) and short-axis (width). The clusters were treated as bivariate normal distributions where we calculated the FWHM of each cluster. For clathrin clusters, the short-axis FWHM is reported.

## Acknowledgements

J.L.H acknowledges funding from the Kathy Chen Fellowship at Columbia University. We acknowledge the Precision Biomolecular Characterization Facility for use of the Zeiss Elyra 7 microscope.

## Code and Data Availability

DBOpt code and simulated datasets are available at https://github.com/Kaufman-Lab-Columbia/DBOpt. k-DBCV code is available at https://github.com/Kaufman-Lab-Columbia/k-DBCV. Cluster simulation code (ClustSim) is available at https://github.com/Kaufman-Lab-Columbia/ClustSim. Experimental data is available upon request.

