## Supporting Information for "Density-based optimization for unbiased, reproducible clustering applied to single molecule localization microscopy"

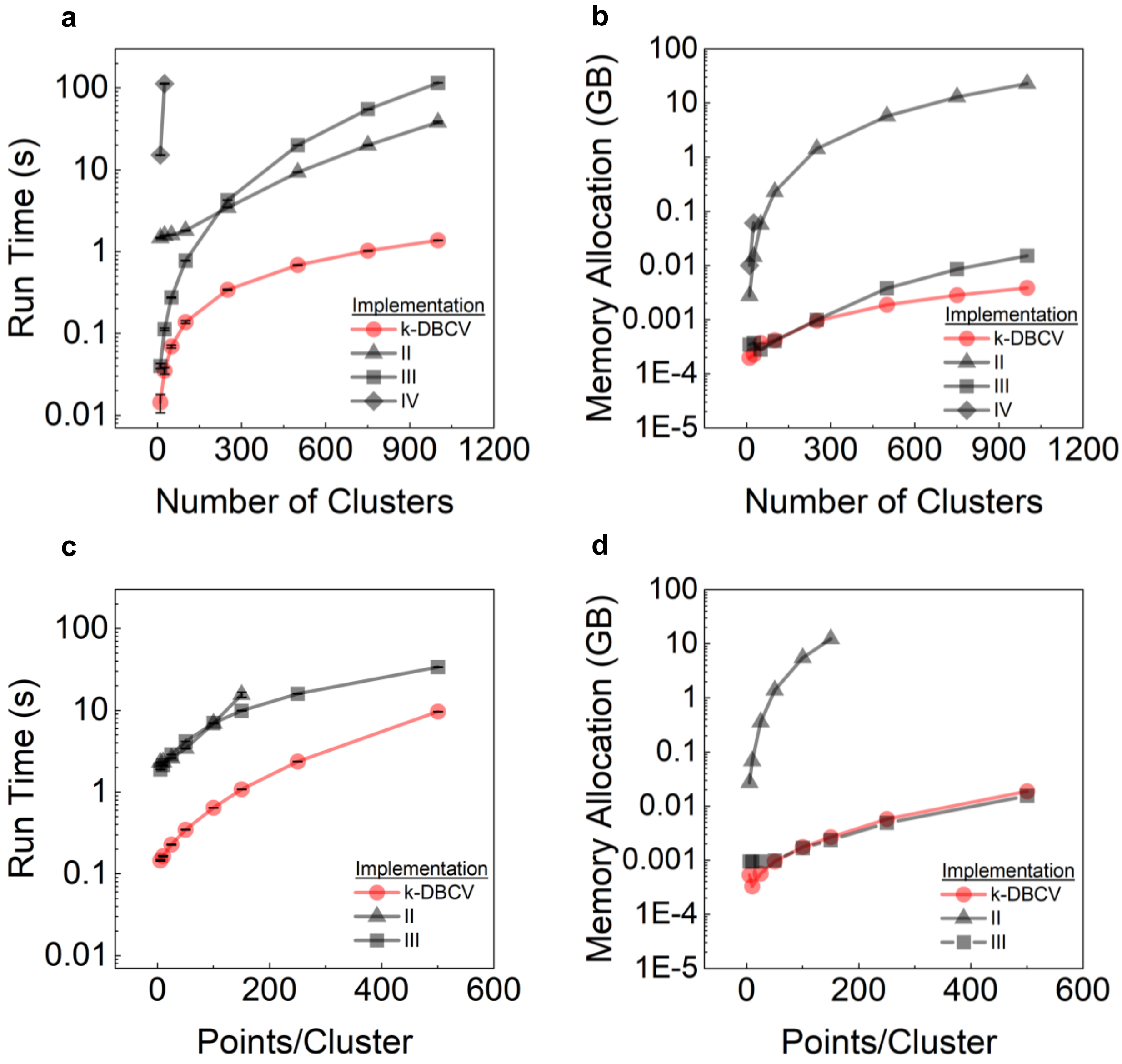


**Fig. S1.** Performance comparisons of the k-DBCV implementation to previous DBCV implementations II[1], III[2], and IV[3]. **a)** Average run time (10 runs, error bars depict standard deviation) and **b)** memory allocation as a function of the number of simulated clusters (each containing 50 points). For implementation IV, data is excluded for cases where run time exceeded 600 s. **c)** Average run time (10 runs, error bars depict standard deviation) and **d)** memory allocation as a function of points per cluster (250 simulated clusters). For implementation IV, data is excluded for cases where run time exceeded 600 s and for implementation II, data is excluded when memory allocation was > 32 GB.

**Supporting Text 1: DBOpt Hyperparameter Selection**

While DBSCAN, HDBSCAN, and OPTICS all have unique input parameters, all share *MinPts* as a common parameter. When choosing the *MinPts* bounds, we suggest a lower bound of $d+1$, where $d$ is the dimensionality of the data, which avoids the commonly described single-link effect.[4] We note that k-DBCV requires a minimum of 3 points to define a cluster, thus in one dimension the *MinPts* should be set to at least 3. While this suggestion serves as an absolute minimum, it is commonly suggested to set the *MinPts* parameter to a value no lower than $2d$.[5] The upper bound of *MinPts* should be set to a number much greater than the expected points per cluster.

The suggested bounds for the remaining algorithm-specific parameters are as follows: for DBSCAN, the lower bound of the $\varepsilon$ parameter should be set near or below the uncertainty of the localizations while the upper bound should be much greater than expected cluster size. For HDBSCAN, *Minimum Cluster Size* bounds should initially be set following the guidance for *MinPts*. For OPTICS, $\xi$ can be set between 0 and 1 to explore the full range of the parameter.

The number of Bayesian iterations for DBOpt to find the parameters that maximize the DBCV score must also be selected. This includes two hyperparameters: the number of parameter combinations to initially randomly seed the parameter space and the number of optimization iterations to sequentially maximize the DBCV score. The sensitivity to the number of optimization iterations is shown in **Fig. S2**, where scores typically converge after 100 optimization iterations. Thus, we suggest a minimum of 100 optimization iterations with the number of parameter combinations to initially seed the parameter space set to 20% of the number of optimization iterations.

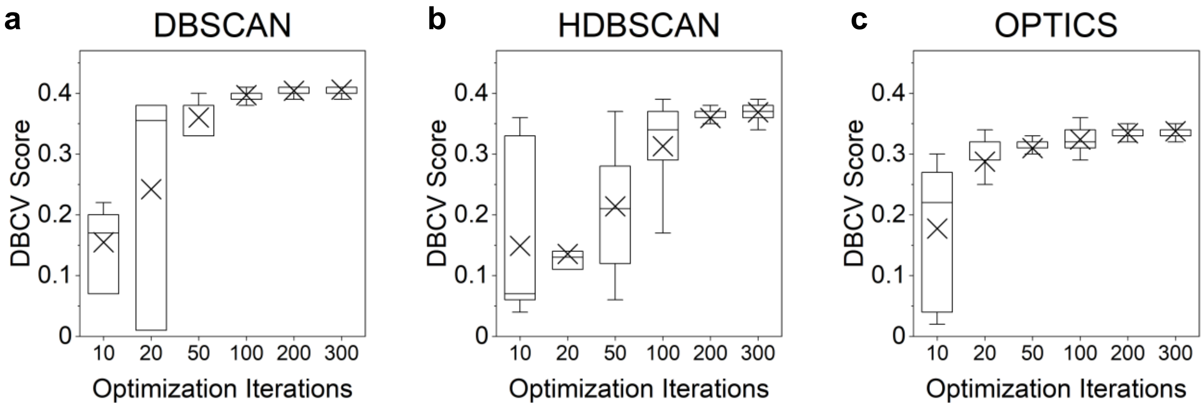


**Fig. S2.** Optimization iteration sweep for **a)** DBSCAN, **b)** HDBSCAN, and **c)** OPTICS where DBOpt is performed on a single simulated dataset (E14 in Table S1) with a set number of optimization iterations preceded by an initial random sampling of parameter combinations (twenty percent of the optimization iterations). Box plot lines indicate median, "X" indicates mean, boxes represent 25$^{th}$ and 75$^{th}$ percentile, and whiskers represent 5$^{th}$ and 95$^{th}$ percentile (n = 50).

### Supporting Text 2: Simulated Clusters

**Table S1a - e** depict ClustSim parameters chosen for simulated circular, elliptic, micellular, fibrillar, and mixed cluster scenarios (ClustSim: https://github.com/Kaufman-Lab-Columbia/ClustSim). Each simulation is named as shown in Column 1, with these names used throughout the manuscript to identify particular simulations. The number of clusters (N) in the simulation and the points per cluster are shown in Columns 2 and 3, respectively. Column 4 shows the width of the short-axis of the cluster for all simulation types, with width reported as 4σ of a normal distribution for circular, elliptic, and fibrillar clusters. For micellular clusters, the width is the outer diameter of the micelle. For circular, elliptic, and micellular clusters, the aspect ratio is the multiple of the width that characterizes the long-axis dimension of the cluster. Aspect ratios depicted as ranges signify an independent, randomly selected aspect ratio for each cluster within the range. Fibrillar cluster length corresponds to the contour length of each fiber with ranges indicating independent, randomly selected lengths for each cluster within the range. The number of noise points in each simulation is reported in Column 5. The user defined noise points, number of clusters, and points per cluster determine the corresponding noise percentage, which is defined as number of noise points divided by total points. 3D spherical cluster simulation parameters are shown in **Table S2**. Here, all parameters can be interpreted as described for 2D circular simulations.

**Table S1a. Circular Simulations**

| Sim. | N (clusters) | Pts/cluster | Width (nm) | Aspect Ratio | Noise pts | % Noise |
|---|---|---|---|---|---|---|
| C01 | 80 | 25 | 100 | 1 | 3000 | 60.0 |
| C02(X)+++ | 80 | 25 | 100 | 1 | 3000* | 60.0 |
| C03 | 50 | 20 | 100 | 1 | 1500* | 60.0 |
| C04 | 50 | 20 | 100 | 1 | 1500 | 60.0 |
| C05 | 50 | 10 | 100 | 1 | 500* | 50.0 |
| C06 | 50 | 50 | 100 | 1 | 1500 | 37.5 |
| C07 | 20 | 100 | 100 | 1 | 1500 | 42.9 |
| C08 | 20 | 10 | 70 | 1 | 0 | 0.0 |
| C09 | 15 | 50 | 300 | 1 | 1200 | 61.5 |
| C10 | 15 | 50 | 300 | 1 | 500 | 40.0 |
| C11 | 85 | 35 | 120 | 1 | 5000* | 62.7 |
| C12 | 10 | 50 | 100 | 1 | 750 | 60.0 |
| C13 | 30 | 30 | 200 | 1 | 1000* | 52.6 |
| C14 | 50 | 25 | 100 | 1 | 500 | 28.6 |
| C15 | 200 | 12 | 50 | 1 | 3000 | 55.6 |
| C16 | 20 | 500 | 200 | 1 | 10000 | 50.0 |
| C17 | 100 | 50 | 75 | 1 | 1000* | 16.7 |
| C18 | 50 | 50 | 100 | 1 | 0 | 0.0 |
| C19+++ | 50 | 15 | 75 | 1 | 1000 | 57.1 |
| C20 | 50 | 30 | 100 | 1 | 1000* | 40.0 |
| C21 | 50 | 10-50 | 100 | 1 | 1500 | 48.2 |
| C22 | 40 | 20-70 | 30-200 | 1 | 1500 | 38.5 |
| C23 | 5 | 30 | 100 | 1 | 200 | 57.1 |
| C24 | 50 | 20-150 | 20-250 | 1 | 5000* | 55.9 |
| C25 | 30 | 10-50 | 100 | 1 | 0 | 0.0 |

(X) denotes a representative simulation shown in Fig. 1 and Fig. 2.
* denotes gradient noise
+++ denotes a corresponding multi-emitter simulation.

**Table S1b. Elliptic Simulations**

| Sim. | N (clusters) | Pts/cluster | Width (nm) | Aspect Ratio | Noise pts | % Noise |
|---|---|---|---|---|---|---|
| E01 | 25 | 100 | 100 | 1-4 | 3000 | 54.5 |
| E02 | 70 | 50 | 100 | 1-4 | 3000 | 46.2 |
| E03 | 40 | 15 | 75 | 1-4 | 1000* | 62.5 |
| E04+++ | 25 | 50 | 100 | 1-4 | 1500 | 54.5 |
| E05 | 40 | 20-70 | 100 | 1-4 | 3000* | 64.8 |
| E06 | 100 | 15 | 50 | 1-4 | 2000* | 57.1 |
| E07 | 15 | 100 | 250 | 1-4 | 2000 | 57.1 |
| E08 | 30 | 20 | 70 | 1-4 | 1000 | 62.5 |
| E09 | 20 | 50 | 100 | 1-4 | 1500 | 60.0 |
| E10 | 50 | 30 | 100 | 2 | 1500 | 50.0 |
| E11 | 20 | 50 | 70 | 6 | 1500 | 60.0 |
| E12 | 50 | 20 | 100 | 1-4 | 0 | 0.0 |
| E13+++ | 45 | 20 | 80 | 1-4 | 500 | 35.7 |
| E14(X) | 50 | 15-40 | 100 | 1-4 | 1500 | 53.2 |
| E15 | 40 | 20-70 | 30-150 | 1-4 | 1500* | 52.9 |
| E16 | 5 | 30 | 100 | 1-4 | 200 | 57.1 |
| E17 | 50 | 35-80 | 30-200 | 1-4 | 4000* | 58.9 |
| E18 | 30 | 10-50 | 100 | 1-4 | 0 | 0.0 |
| E19 | 200 | 5 | 20 | 1-4 | 1000 | 50.0 |
| E20 | 30 | 18 | 120 | 1-4 | 300* | 35.7 |
| E21 | 50 | 15 | 80 | 1-2 | 1000 | 57.1 |
| E22 | 25 | 30 | 80 | 1-6 | 1000 | 57.1 |
| E23 | 12 | 50 | 150 | 4 | 1000 | 62.5 |
| E24 | 25 | 20 | 100 | 1-4 | 0 | 0.0 |
| E25 | 30 | 15-40 | 100 | 1-4 | 1100* | 58.9 |

(X) denotes a representative simulation shown in Fig. 1 and Fig. 2.
* denotes gradient noise
+++ denotes a corresponding multi-emitter simulation.

**Table S1c. Micellular Simulations**

| Sim. | N (clusters) | Pts/cluster | Width (nm) | Aspect Ratio | Noise pts | % Noise |
|---|---|---|---|---|---|---|
| M01 | 30 | 50 | 150 | 1 | 2000* | 57.1 |
| M02 | 80 | 50 | 150 | 1 | 5000 | 55.6 |
| M03 | 50 | 20 | 100 | 1 | 1200 | 54.5 |
| M04 | 70 | 23 | 100 | 1 | 3000 | 65.1 |
| M05 | 30 | 25 | 100 | 1 | 0 | 0.0 |
| M06+++ | 30 | 45 | 100 | 1 | 1500 | 52.6 |
| M07 | 50 | 200 | 200 | 1 | 7000* | 41.2 |
| M08 | 50 | 60 | 100 | 1-3 | 3000* | 50.0 |
| M09+++ | 40 | 60 | 150 | 1 | 3000* | 55.6 |
| M10 | 50 | 40-80 | 80-150 | 1 | 3500 | 54.0 |
| M11 | 28 | 55-110 | 120 | 1 | 3500* | 60.8 |
| M12 | 15 | 20 | 80 | 1 | 500* | 62.5 |
| M13 | 100 | 20 | 80 | 1 | 0 | 0.0 |
| M14 | 60 | 40 | 130 | 1 | 500 | 17.2 |
| M15 | 10 | 150 | 300 | 1 | 1500 | 33.3 |
| M16 | 30 | 60-100 | 180-240 | 1 | 2500* | 51.0 |
| M17 | 50 | 35 | 80 | 2 | 1200 | 40.7 |
| M18 | 40 | 50-75 | 80-120 | 1-2 | 3000* | 54.1 |
| M19 | 20 | 30 | 100 | 1 | 200 | 25.0 |
| M20 | 5 | 30 | 100 | 1 | 200 | 57.1 |
| M21 | 40 | 55 | 120-160 | 1 | 3000 | 57.7 |
| M22(X) | 30 | 100 | 200 | 1 | 2500 | 45.5 |
| M23 | 20 | 30-50 | 80-140 | 1 | 1200* | 60.1 |
| M24 | 20 | 20 | 100 | 1 | 100 | 20.0 |
| M25 | 200 | 20 | 80 | 1 | 3000* | 42.9 |

(X) denotes a representative simulation shown in Fig. 1 and Fig. 2.
* denotes gradient noise
+++ denotes a corresponding multi-emitter simulation.

**Table S1d. Fibrillar Simulations**

| Sim. | N (clusters) | Pts/cluster | Width (nm) | Length (nm) | Noise pts | % Noise |
|---|---|---|---|---|---|---|
| F01 | 6 | 500 | 50 | 1000 | 3000 | 50.0 |
| F02 | 6 | 3000 | 50 | 1000 | 3000 | 14.3 |
| F03 | 6 | 1000 | 50 | 1000 | 3000 | 33.3 |
| F04 | 3 | 1000 | 50 | 2000 | 3000 | 50.0 |
| F05 | 15 | 200 | 50 | 500 | 4000* | 57.1 |
| F06 | 15 | 200 | 50 | 500 | 4000 | 57.1 |
| F07 | 14 | 80 | 100 | 700 | 2000* | 67.1 |
| F08 | 10 | 150 | 100 | 700 | 0 | 0.0 |
| F09 | 12 | 135-200 | 100 | 300-1000 | 3000* | 60.0 |
| F10 | 6 | 250-650 | 50-150 | 300-1000 | 5000* | 49.8 |
| F11 | 3 | 700 | 100 | 1000 | 3000* | 58.8 |
| F12+++ | 20 | 50 | 50 | 500 | 1000 | 50.0 |
| F13 | 12 | 200 | 50 | 500 | 800 | 25.0 |
| F14+++ | 20 | 150 | 50 | 500 | 4000* | 57.1 |
| F15 | 5 | 1000 | 150 | 1000 | 4000 | 44.4 |
| F16 | 5 | 1000 | 25 | 1000 | 7500 | 60.0 |
| F17 | 15 | 260-320 | 120 | 500-1000 | 5000* | 53.5 |
| F18(X) | 18 | 110-170 | 70 | 300-800 | 3000* | 54.3 |
| F19 | 10 | 200 | 70 | 800 | 3000 | 65.2 |
| F20 | 3 | 2000 | 120 | 2000 | 0 | 0.0 |
| F21 | 12 | 500 | 100 | 500 | 5000 | 45.5 |
| F22 | 4 | 500 | 100 | 500 | 3000 | 60.0 |
| F23 | 7 | 80 | 150 | 1000 | 300 | 30.0 |
| F24 | 5 | 1000 | 25 | 1000 | 2000 | 28.6 |
| F25 | 12 | 150 | 50 | 700 | 2000* | 54.3 |

(X) denotes a representative simulation shown in Fig. 1 and Fig. 2.

* denotes gradient noise

+++ denotes a corresponding multi-emitter simulation.

**Table S1e. Mixed Simulations**

| Sim. | Cluster Type | N (clusters) | Pts / cluster | Width (nm) | Aspect Ratio / (Length (nm)) | Noise pts | % Noise |
|---|---|---|---|---|---|---|---|
| V01 | E, F | 10,5 | 50, 500 | 100, 50 | 1-4, (700) | 3000 | 50.4 |
| V02 | C, F | 10,5 | 50, 500 | 100, 50 | 1, (700) | 3000 | 50.4 |
| V03 | M, F | 10,5 | 100, 500 | 200, 50 | 1, (700) | 3000 | 46.5 |
| V04 | E, M | 10,10 | 60, 120 | 100, 200 | 1-4, 1 | 2500 | 58.1 |
| V05 | E, M | 10,12 | 50, 100 | 100, 150 | 1-4, 1 | 2500* | 59.5 |
| V06 | E, M, F | 7,7,5 | 50, 100,500 | 100, 150, 50 | 1-4, 1, (500) | 2500* | 41.3 |
| V07 | E, M, F | 7,7,5 | 50, 100,500 | 100, 150, 50 | 1-4, 1, (500) | 0 | 0.0 |
| V08 | E, M | 12,12 | 50, 100 | 100, 150 | 1-4, 1 | 0 | 0.0 |
| V09+++ | E, F | 15,7 | 40, 200 | 70, 50 | 1-4, (250) | 1500* | 42.9 |
| V10 | E, M, F | 7,7,6 | 50, 100,300 | 100, 200, 50 | 1-4, 1, (700) | 3000* | 51.3 |
| V11+++ | M, F | 5,9 | 100, 200 | 100, 100 | 1-4, (500) | 2000 | 46.5 |
| V12(X) | C, E, M, F | 10,6,6,9 | 30, 60, 150, 200 | 70, 100, 150, 50 | 1, 1-4, 1, (500) | 3000* | 47.2 |
| V13 | C, M | 12,15 | 30, 100 | 70, 150 | 1, 1 | 800 | 30.1 |
| V14 | C, F | 12,7 | 30, 150 | 70, 70 | 1, (600) | 1200 | 50.0 |
| V15 | E, F | 10,8 | 45, 150 | 120, 30 | 1-4, (600) | 900 | 46.0 |
| V16 | C, E | 15,10 | 20, 50 | 30, 80 | 1 | 700 | 46.7 |
| V17 | E, M, F | 6,6,4 | 60, 70, 200 | 60, 120, 40 | 1-4, 1, (500) | 2000 | 47.8 |
| V18 | E, F | 15,5 | 60, 200 | 100, 50 | 1-4, (600) | 0 | 0.0 |
| V19 | M, F | 15,5 | 80, 200 | 120, 50 | 1, (600) | 0 | 0.0 |
| V20 | C, F | 15,3 | 50, 400 | 100, 100 | 1, (1600) | 3000* | 63.7 |
| V21 | C, E, M, F | 7,7,7,6 | 50, 60, 80, 120 | 80, 100, 120, 40 | 1, 1-4, 1, (400) | 2000 | 49.4 |
| V22 | C, E, M, F | 7,7,7,6 | 50, 60, 80, 120 | 80, 100, 120, 40 | 1, 1-4, 1, (400) | 0 | 0.0 |
| V23 | C, E, F | 9,9,7 | 50, 60, 150 | 80, 100, 60 | 1, 1-4, (500) | 3000* | 59.5 |
| V24 | E, M, F | 9,9,6 | 50, 130, 150 | 80, 100, 60 | 1-4, 1-3, (500) | 1200* | 32.3 |
| V25 | C, E, M | 12,12,12 | 40, 60, 100 | 60, 100, 120 | 1, 1-4, 1 | 2500* | 51.0 |

(X) denotes a representative simulation shown in Fig. 1 and Fig. 2
C, E, M, and F denote Circular, Elliptic, Micellular, and Fibrillar clusters, respectively
* denotes gradient noise
+++ denotes a corresponding multi-emitter simulation.

**Table S2. 3D Simulations**

| Sim. | N (clusters) | Pts/cluster | Width (nm) | Aspect Ratio | Noise pts | % Noise |
|---|---|---|---|---|---|---|
| S01(X) | 50 | 50 | 100 | 1 | 3000 | 54.5 |
| S02 | 50 | 50 | 100 | 1 | 1500 | 37.5 |
| S03 | 50 | 50 | 100 | 1 | 500 | 16.7 |
| S04 | 50 | 50 | 100 | 1 | 0 | 0.0 |
| S05 | 50 | 25 | 100 | 1 | 1500 | 54.5 |
| S06 | 50 | 100 | 100 | 1 | 1500 | 23.1 |
| S07 | 50 | 10 | 40 | 1 | 500 | 50.0 |
| S08 | 50 | 25 | 200 | 1 | 1500 | 54.5 |
| S09 | 25 | 50 | 200 | 1 | 1500 | 54.5 |
| S10 | 100 | 10 | 40 | 1 | 1500 | 60.0 |

(X) denotes a representative simulation shown in Fig. 4.

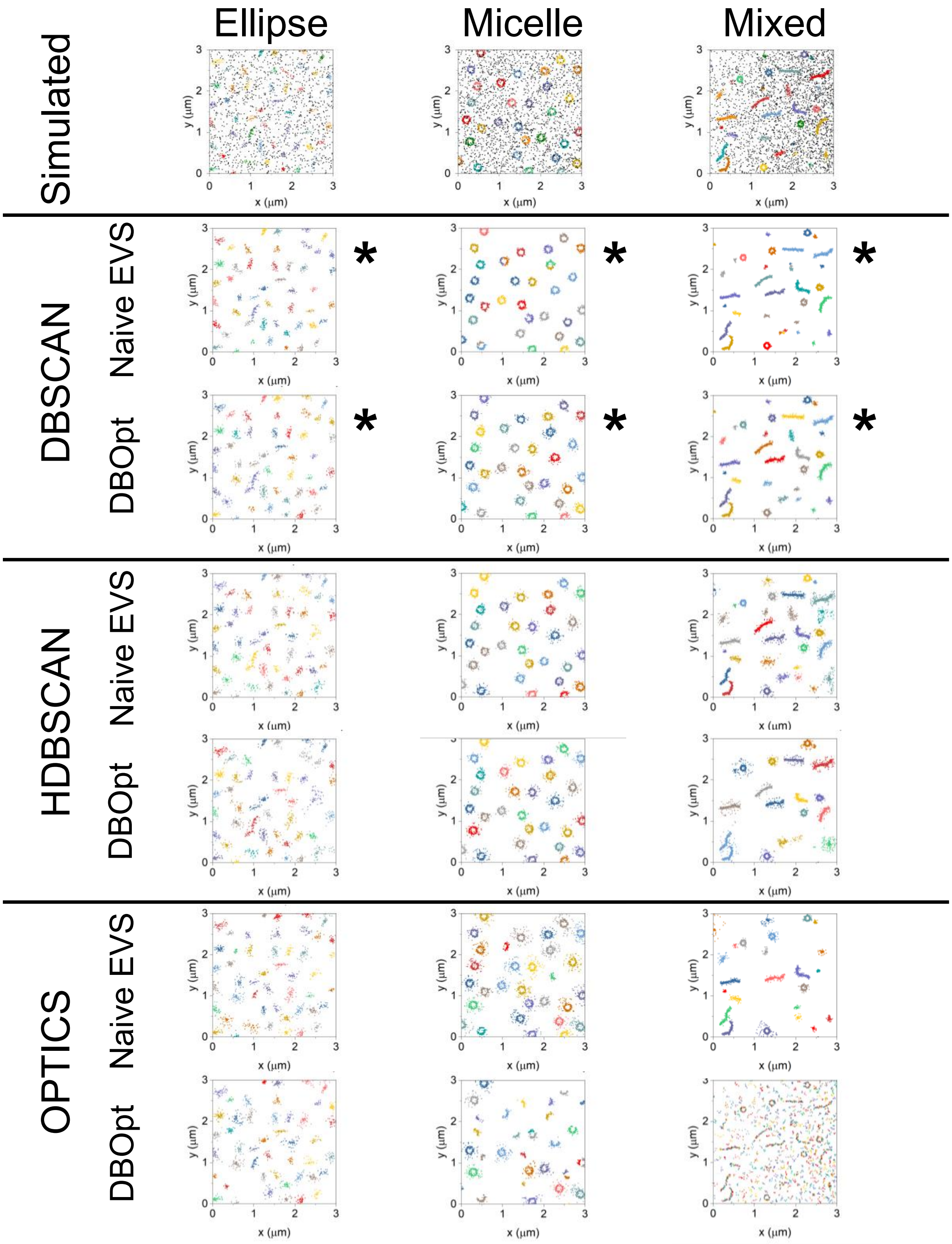


**Fig. S3.** Representative plots for simulated ellipse (E14), micelle (M22), and mixed (V12) clustering scenarios depicting the clustering results with noise removed for visualization for naive EVS and DBOpt for DBSCAN, HDBSCAN, and OPTICS. * indicates best performing algorithm for EVS or DBOpt. Selected plots are depicted with noise in **Fig. 2** in the main text.

**Supporting Text 3: Multi-emitters**

In SMLM, multiple localizations associated with the same target molecule are common, as many SMLM approaches employ antibodies that have multiple fluorophores per protein.[6] To improve the correspondence of simulated datasets to experiments, we simulated multi-emitter 2D data where the number of localizations per molecule was drawn from a Poisson distribution with the average localizations per simulated molecule set to 3.[7] We then compared the performance of DBOpt on multi-emitter data to the previously simulated, single localization data (**Fig. S4**). While a slight decrease in performance for DBSCAN and HDBSCAN is observed using the original hyperparameters, increasing the lower bound of *MinPts* from 3 to 6 ($2(d+1)$), corresponding to the assumption that the average molecule will be represented by at least two localizations, fully recovers and in some cases improves performance relative to single-emitter data (**Fig. S4c**).

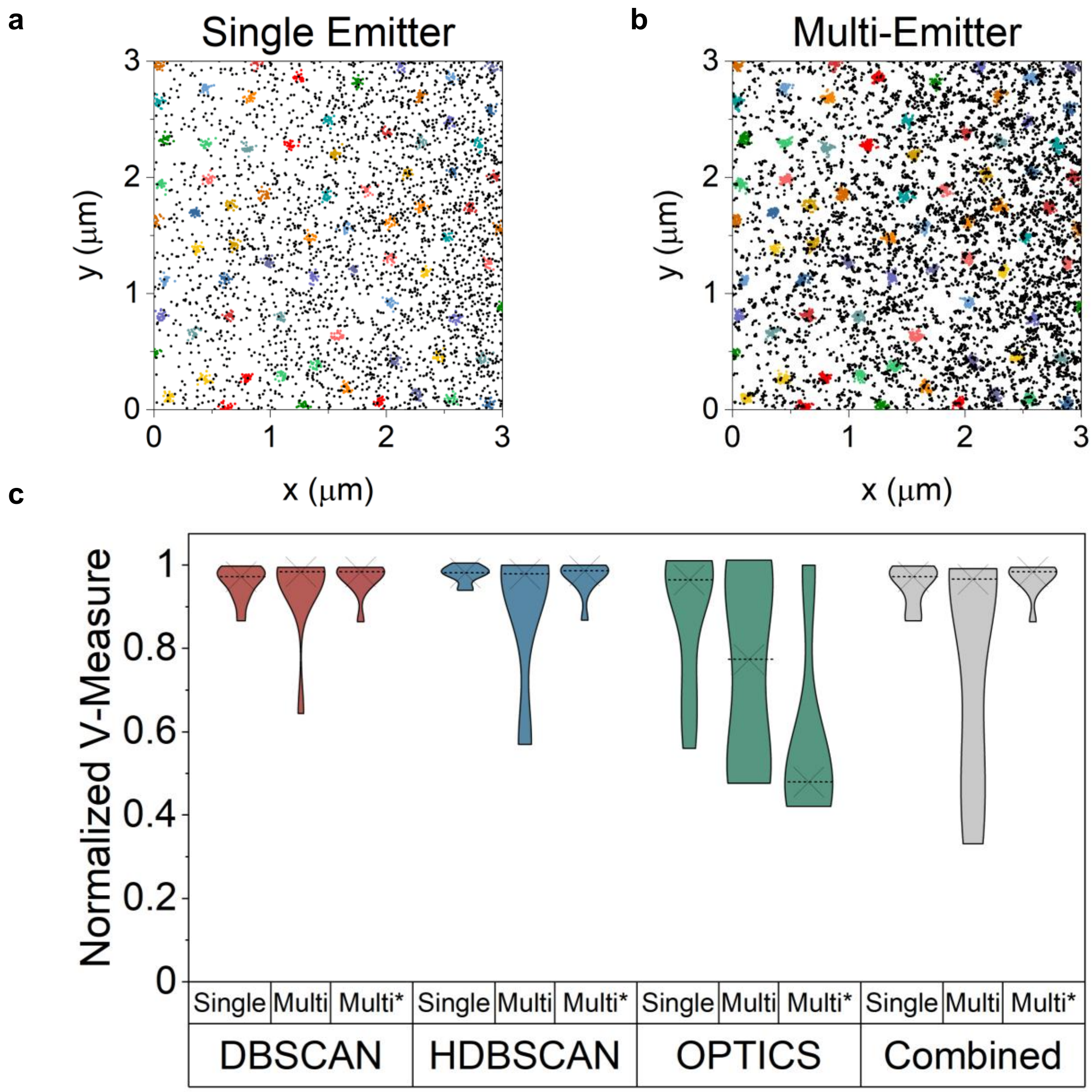


**Fig. S4.** Representative simulation (C02) of **a)** single emitter and **b)** multi-emitter data. **c)** Comparisons of normalized V-measure score distributions for ten simulations where the ratios of DBOpt to naive EVS V-measure scores were calculated for corresponding single multi-emitter datasets (dotted line represents median and "X" represents mean). Multi* indicates evaluation in which the lower bound of *MinPts* was set to of 6. Simulations are denoted in Tables S1a-e by $^{+++}$.

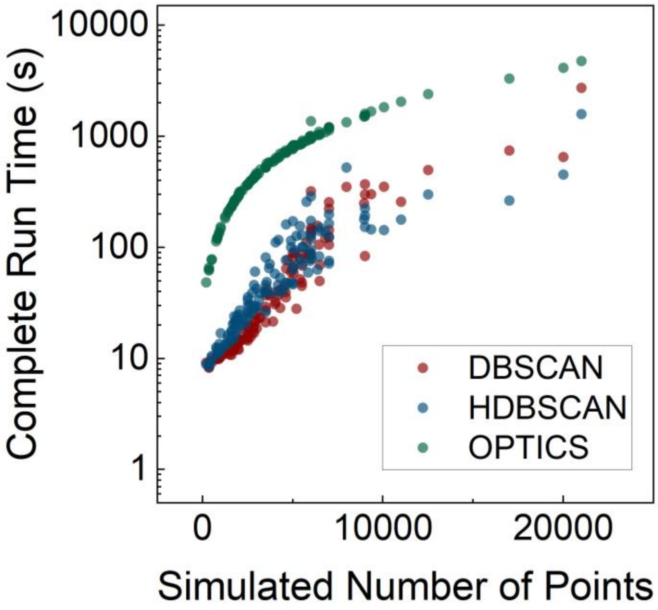


**Fig. S5.** Full runtime of DBOpt with 40 initial random parameter sets and 200 subsequent optimization iterations for DBSCAN, HDBSCAN, and OPTICS on all 2D single-emitter simulated datasets listed in Table S1a-e.

**Supporting Text 4: Noise Evaluation**

To assess the sensitivity of DBOpt to noise, evaluation of DBOpt performance on randomly generated noise in the absence of clusters was performed. Because noise points will not be completely homogeneous, a positive DBCV score for simulated SMLM noise clustered with DBOpt-selected parameters was expected. We generated ten simulations at multiple point densities and performed DBOpt, varying the lower bound of the *MinPts* parameter for each algorithm, as this parameter was expected to have the greatest impact on each algorithm's tendency to cluster noise. We show the maximum DBCV scores from each set of ten simulations in **Fig. S6** for 2D and 3D simulations. The score is dependent on the *MinPts* parameter, noise density, and the number of dimensions. These scores should be kept in mind when evaluating experimental data, as when the DBOpt result is less than the corresponding noise scores, it is not possible to determine if DBOpt is clustering data distinct from noise.

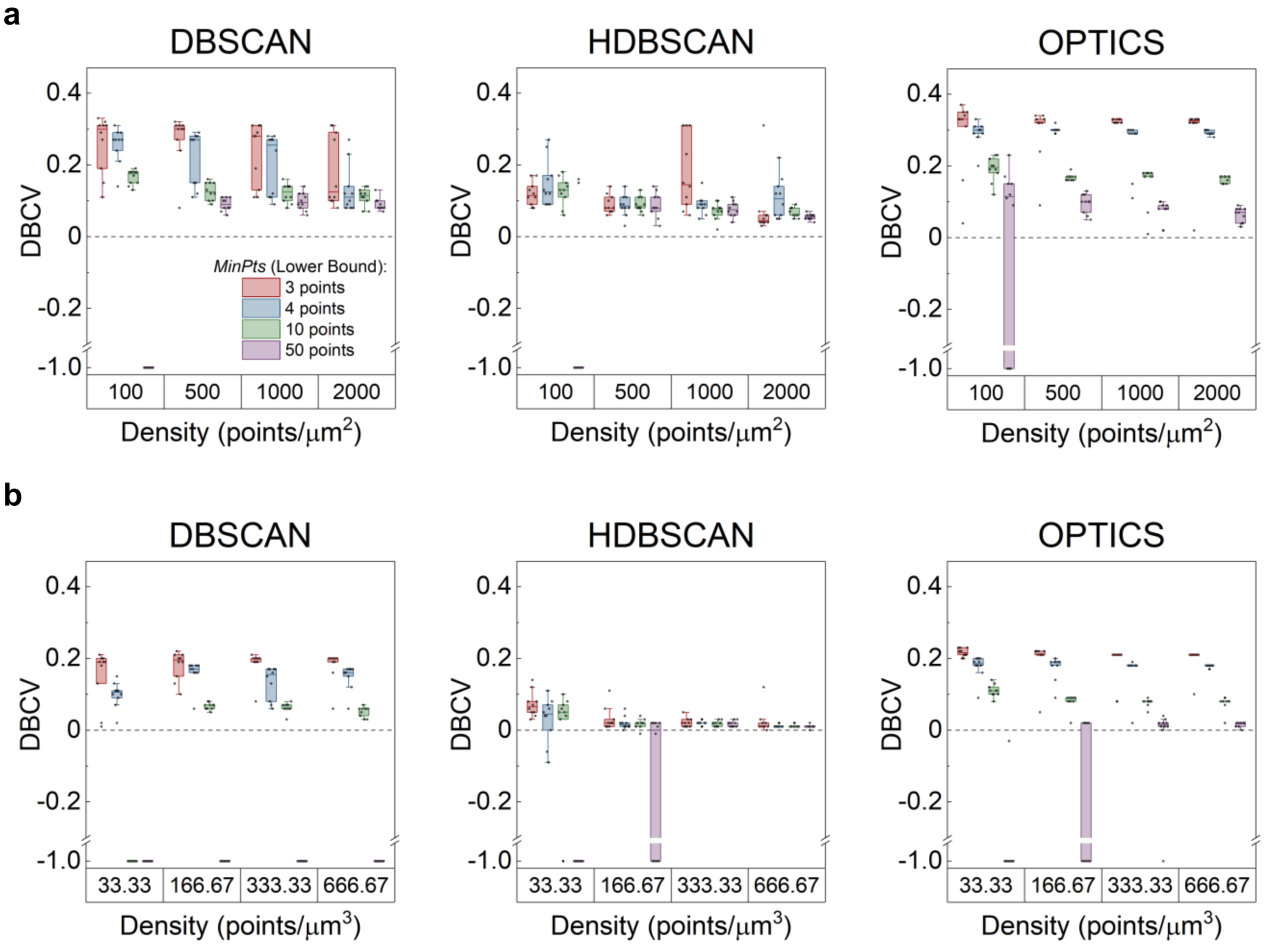


**Fig. S6.** DBOpt DBCV scores for n = 10 noise simulations for DBSCAN, HDBSCAN, and OPTICS in **a)** 2D and **b)** 3D, varying density and the lower bound of the *MinPts* parameter. Box plot lines indicate median, boxes represent 25$^{th}$ and 75$^{th}$ percentile, and whiskers represent 5$^{th}$ and 95$^{th}$ percentile. Legend in 2D DBSCAN plot applies to all plots.

**Supporting Text 5: Experimental Data**

Experimental data for integrin and clathrin is described in **Table S3**. The dataset name is shown in Column 1, where each dataset corresponds to an image of a single cell. After clustering data from the full cell, the DBCV score and optimal parameters from DBOpt (paired with DBSCAN) are shown in Columns 2-4, respectively. Noise thresholds shown in Column 5 are approximated from **Fig. S6**. Here, the maximum DBCV score for each *MinPts* parameter is determined and set as the noise threshold. When the experimental *MinPts* parameter is not equal to any of the values shown in **Fig. S6**, the noise threshold ($T_n$) is expected to be between the thresholds of the noise-only simulations using the nearest evaluated *MinPts* parameters. In some cases, the noise threshold is found to be -1. This indicates that DBOpt paired with DBSCAN is unable to find a set of parameters for the simulated noise scenario that identifies two or more clusters and no DBCV score can be calculated. Thus, while no noise threshold is relevant for these scenarios, reverting to a threshold of 0 should be considered best practice as negative DBCV scores signify data where sparseness is greater than separation, which is an unphysical clustering result. In all cases the DBCV score is above the maximum score associated with noise alone, validating the clustering achieved in this experimental data.

**Table S3. Experimental Clustering Summary**

| Dataset | DBCV Score | ε | *MinPts* | Noise Threshold (Fig. S6) |
|---|---|---|---|---|
| Integrin 01 | 0.59 | 63.20 | 7 | $0.19 < T_n < 0.31$ |
| Integrin 02 (X) | 0.55 | 38.24 | 6 | $0.19 < T_n < 0.31$ |
| Integrin 03 | 0.53 | 47.13 | 13 | $0.14 < T_n < 0.19$ |
| Integrin 04 | 0.63 | 63.29 | 4 | 0.31 |
| Integrin 05 | 0.61 | 55.05 | 3 | 0.33 |
| Clathrin 01 (X) | 0.12 | 126.80 | 46 | $-1 < T_n < 0.08$ |
| Clathrin 02 | 0.12 | 111.66 | 34 | $-1 < T_n < 0.08$ |
| Clathrin 03 | 0.15 | 132.11 | 52 | -1 |
| Clathrin 04 | 0.09 | 135.69 | 60 | -1 |
| Clathrin 05 | 0.10 | 196.99 | 112 | -1 |

(X) denotes experimental data shown in Fig. 5.
